# High-throughput phenotyping of soybean transpiration response curves to rising atmospheric drying in a mapping population

**DOI:** 10.1101/2024.05.16.594513

**Authors:** Daniel Monnens, Erik McCoy, Bishal G. Tamang, Aaron J. Lorenz, Walid Sadok

## Abstract

In soybean, limiting whole-plant transpiration rate (TR) response to increasing vapor pressure deficit (VPD) has been associated with the ‘slow-wilting’ phenotype and with water-conservation enabling higher yields under terminal drought. Despite the promise of this trait, it is still unknown whether it has a genetic basis in soybean, a challenge limiting the prospects of breeding climate-resilient varieties. Here we present the results of a first attempt at a high-throughput phenotyping of TR and stomatal conductance response curves to increasing VPD conducted on a soybean mapping population consisting of 140 recombinant inbred lines (RIL). This effort was conducted over two consecutive years, using a controlled-environment, gravimetric phenotyping platform which enabled characterizing 900 plants for these responses, yielding regression parameters (R^2^ from 0.92 to 0.99) that were used for genetic mapping. Several quantitative trait loci (QTL) were identified for these parameters on chromosomes (Ch) 4, 6 and 10, including a VPD-conditional QTL on Ch 4 and a ‘constitutive’ QTL controlling all parameters on Ch 6. This study demonstrated for the first time that canopy water use in response to rising VPD has a genetic basis in soybean, opening novel avenues for identifying alleles enabling water conservation under current and future climate scenarios.

## Introduction

In many agricultural hotspots across the globe, climate change is reducing crop productivity, largely through effects that alter crop water balance (e.g., Lobell *et al*. 2014; Zhang *et al*. 2017; Kimm *et al*. 2020). A key, yet poorly understood driver of climate change-driven water deficits is atmospheric water vapor pressure deficit (VPD), an environmental variable that drives evaporation from soil and plant canopies (Grossiord *et al*. 2020; López *et al*. 2021).

Recently, it has been observed that a worldwide increase in VPD is taking place and that this increase is expected to continue to amplify over the next decades, negatively impacting yields (Ficklin and Novick, 2017; Yuan *et al*. 2019; Sun *et al*. 2023).

In soybean, continent-wide increases in seasonal VPD observed across 27 U.S. states from 2007 to 2016 have been shown to be the most important drivers of yield variation, and this effect was detected even in the typically well-watered Upper Midwest (Mourtzinis *et al*. 2019). The effects of VPD on plant physiology and productivity are complex and still incompletely understood (Sinclair *et al*. 2017; Grossiord *et al*. 2020; López *et al*. 2021). However, it is widely accepted that negative effects on soybean yields arising from VPD increase are at least partly driven by higher crop water needs due to transpiration rate (TR) increase (Fletcher *et al*. 2007; Sinclair *et al*. 2010). Because crop productivity is largely constrained by seasonal water availability, increases in TR that are not matched by adequate water supply typically lead to exposing flowering and seed-fill to higher levels of soil moisture deficit, that is, terminal drought. Due to the critical importance of reproductive development in determining final yield, this pattern of water deficit typically translates into to yield penalties (Sinclair *et al*. 2005; Messina *et al*. 2015; Sadok *et al*. 2019).

In soybean, ‘slow-wilting’ genotypes, which exhibit superior drought tolerance under terminal drought conditions, were found to exhibit a limitation in their TR when VPD exceeded a threshold of 1.5-2 kPa, which typically occurs around midday (Fletcher *et al*. 2007; Sadok and Sinclair 2009ab; Sadok and Sinclair 2010a). Simulation modelling indicated that this water conservation strategy enables higher levels of stored soil moisture during seed fill, thereby leading to yield gains across a range of production environments in the U.S. (Sinclair *et al*. 2010). This behaviour was first reported in a soybean line PI 146937 (Fletcher *et al*. 2007), which has been since then successfully used a donor line to various breeding programs across the U.S. (Carter *et al*. 2016; Ye *et al*. 2020). The physiological basis of this water-saving behaviour was shown to be driven by a hydraulic limitation of water transport in the leaves between the xylem and the guard cells (Sinclair *et al*. 2008). This hydraulic restriction has subsequently been linked to a lack of silver-sensitive aquaporin population that mediates cell-to-cell water movement in the mesophyll of the leaves of the slow-wilting genotype PI 416937 (Sadok and Sinclair 2010b).

Using this information, efforts have been undertaken to screen mapping populations by feeding de-rooted soybean plants with silver nitrate, a compound known to inhibit silver- sensitive aquaporins, as an indirect way to identify the genetic basis of the water-conservation trait (Carpentieri-Pipolo *et al*. 2012; Steketee *et al*. 2019; Sarkar *et al*. 2020). However, a major challenge with these approaches, remains that they are fundamentally based on using toxic chemicals applied on non-intact plants and as a result, they may capture a host of physiological side-effects that could generate artefactual trait-marker associations.

Directly phenotyping mapping populations for their TR response to VPD is a notoriously challenging effort and has not been successfully conducted on soybean. Since 2007, the cumulative number of soybean genotypes that were screened for their whole-plant TR response to VPD is probably under 60 (Fletcher *et al*. 2007; Sinclair *et al*. 2008; Sadok and Sinclair 2009ab; Seversike *et al*. 2014; Devi *et al*. 2014). This is mainly due to the complex logistics of i) reliably tracking intact, whole-plant TR, ii) imposing target VPD levels consistently and in a way that enables generating response curves, and iii) minimizing VPD interaction with other potentially confounding environmental variables that are known to influence TR (e.g., temperature, light, watering, air mixing). Due to these limitations, the extent of genetic variability in this trait in soybean and its genetic basis remain unknown.

To address these challenges, we developed a gravimetric screening system to be deployed across three identical walk-in growth chambers equipped with ad hoc VPD control systems. This platform was successfully used to phenotype high-resolution TR response curves to VPD on nested association mapping (NAM) parents in a number of crops, revealing substantial phenotypic diversity in maize (Tamang and Sadok 2018), wheat (Tamang *et al*. 2019), and barley (Sadok and Tamang 2019). More recently, this platform enabled identifying the genetic basis of this trait in wheat (Tamang *et al*. 2022).

The first objective of this research was to characterize, using this phenotyping platform, the extent of genotypic variability in TR response to VPD within a mapping population consisting of 140 recombinant inbred lines (RILs). These lines are derived from a cross between a commercial line, IA3023, and LG94-1906, which were identified based on a pilot experiment as exhibiting relatively contrasting TR response curves to VPD. The second objective of the study was to conduct a genetic mapping approach to identify QTL controlling TR response to VPD in soybean for the first time.

## Materials and methods

### Genetic material

The genetic material of this study consisted of 140 RILs of soybean derived from a cross between parents IA3023 and LG03-3191, a line with diverse ancestry (Diers *et al*. 2018). This population, called NAM25, is part of the soybean nested association mapping panel (SoyNAM), which is composed of 40 soybean varieties crossed to a common parent, IA3023, to develop 40 RIL populations (www.soybase.org/SoyNAM). The NAM25 population was selected based on a preliminary experiment where all 40 parents were screened for their TR response to VPD, in which LG03-3191 exhibited a response that contrasted the most with that of the recurrent parent IA3023 (Tamang *et al*. Unpublished).

### Experimental design and plant growth conditions

The population of 140 RILs and their parents were phenotyped in two independent experiments conducted at St. Paul campus of the University of Minnesota (44.9866° N, 93.1832° W) over 2 consecutive years (2018-2019), using a gravimetric phenotyping platform (GraPh system, Tamang and Sadok 2018; Tamang *et al*. 2019). This system enables the screening of up to 54 plants simultaneously on a given day (18 genotypes replicated three times) under controlled environment conditions. Due to the logistics of the phenotyping (see below for details), this effort took place over 3 consecutive days in a given week, resulting in a throughput of 54 genotypes per week. Therefore, in order to phenotype a population of 140 RILs, three sequential plantings – one per week – were implemented in each experiment, with each group consisting of 45-50 RILs and one of the two parents as checks. The experimental design was an augmented incomplete block design with planting date as block and one of the parental lines as check. Parents LG03-3191 and IA3023 were the checks for experiments 1 and 2 respectively.

Plants were grown for 28-44 days until they reached the V4 stage, in a glasshouse regulated for a minimal temperature of 18°C (Table 1), based on a protocol developed by Tamang *et al*. (2019). Briefly, seeds were planted at a depth of 2.5 cm in 3.8-liter pots filled with a garden mix (35% organic compost, 25% course fibrous peat, 30% screened soil, and 10% course sand; Plaisted Companies Inc., Elkriver, MN, USA). Pot filling was achieved such that seeds were planted in pots filled with a uniform amount of soil. In each pot, three similarly sized seeds were placed into three equidistant holes in the center of the pot, and then covered with powdered rhizobial inoculant of *Bradyrhizobium japonicum* (NDure, Cary, NC). All pots were then lightly watered and immediately covered with tarps to minimize water evaporation from the surface of the soil until seedlings emerged, on average 4 d after planting.

**Table 1.** Growth conditions subjected to the NAM25 population recombinant inbred lines (RILs) during experiments E1 and E2. Averaged (± S.E.) daytime and nighttime conditions are presented for each planting group.

| Exp. | Group | Daytime conditions |  |  |  | Night-time conditions |  |  |  |
| --- | --- | --- | --- | --- | --- | --- | --- | --- | --- |
| | | T (C°)<br>± S.E. | RH (%)<br>± S.E. | VPD<br>(kPa)<br>± S.E. | PAR<br>($\mu\text{mol m}^{-2}$<br>$\text{s}^{-1}$) ± S.E. | T (C°)<br>± S.E. | RH (%)<br>± S.E. | VPD<br>(kPa)<br>± S.E. | PAR<br>($\mu\text{mol m}^{-2}\text{s}^{-1}$) |
| **E1** | 1 | 32.8 ± 0.9 | 39.3 ± 2.1 | 3.3 ± 0.3 | 425 ± 23 | 21.6 ± 0.4 | 66.0 ± 2.1 | 0.9 ± 0.1 | 0 |
| | 2 | 32.2 ± 0.9 | 44.3 ± 2.5 | 3.0 ± 0.3 | 394 ± 26 | 22.2 ± 0.3 | 65.9 ± 0.3 | 0.9 ± 0.1 | 0 |
| | 3 | 32.0 ± 0.9 | 48.6 ± 2.9 | 2.8 ± 0.3 | 385 ± 31 | 22.8 ± 0.4 | 67.6 ± 0.4 | 0.9 ± 0.1 | 0 |
| **E2** | 1 | 23.7 ± 0.3 | 45.7 ± 1.8 | 1.7 ± 0.1 | 238 ± 15 | 18.0 ± 0.2 | 57.0 ± 0.2 | 0.9 ± 0.0 | 0 |
| | 2 | 24.0 ± 0.2 | 48.3 ± 1.6 | 1.6 ± 0.1 | 204 ± 10 | 18.9 ± 0.2 | 56.9 ± 0.2 | 1.0 ± 0.0 | 0 |
| | 3 | 24.1 ± 0.3 | 43.4 ± 1.7 | 1.8 ± 0.1 | 210 ± 11 | 19.6 ± 0.0 | 49.4 ± 0.0 | 1.2 ± 0.0 | 0 |

One week following planting, each pot was given an equal amount of a slow-release fertilizer (15-9-12 NPK, Osmocote Plus, Miracle-Gro, Marysville, OH) along with a granular systemic insecticide (Marathon, OHP Inc., Bluffton, SC) to minimize thrips damage. Ten days after planting, pots were thinned from three to two plants, and three days later they were all thinned to one plant per pot. Plants were watered at least twice weekly depending on plant size and evaporative conditions. On average during the growth period, pots were given 300 ml per irrigation event.

During the growth period, temperature (T), relative humidity (RH) and vapor pressure deficit (VPD) conditions inside the glasshouse were recorded every 5 mins with three portable USB sensors (Model EL-USB-2-LCD, Lascar Electronics, Whiteparish, UK) which were placed at canopy height in three different locations across the glasshouse (Table 1). Similarly, photosynthetically active radiation (PAR) was measured at canopy height using a quantum sensor (model S-LIA-M003, Onset Computer Corporation, Bourne, MA) connected to a data logger (HOBO H21-USB micro station, Onset computer Corporation, Bourne, MA) every 5 minutes (Table 1). Plants were randomized in the glasshouse and rotated twice per week to minimize long-term effects on growth arising from spatial heterogeneity in PAR, T and VPD gradients across the glasshouse.

### Phenotyping transpiration rate response curves to increasing vapor pressure deficit

After growing in the glasshouse, plants were moved inside the GraPh platform following the procedure fully detailed in Tamang *et al*. (2022). The platform consists of 54 balances (Model Fx-3000i, A & D Co. Ltd, Tokyo, Japan) protected from dust, moisture, vibration, and static electricity and connected to dataloggers which record changes in pot mass every minute at a resolution of a 0.01 g. These balances were placed inside 3 adjacent, identical, walk-in programmable growth chambers (Model PGV36, Conviron, Controlled Environments Ltd., Winnipeg, Manitoba, Canada), where target environmental conditions were imposed as reported on Table 2. In these experiments, VPD changes were imposed over 7 consecutive steps where VPD was increased from 1.5 kPa to 3 kPa while temperature and PAR were kept constant at 30°C and approx. 500 μmol m^-2^ s^-1^, respectively (Table 2).

**Table 2.** Temperature (T) and vapor pressure deficit (VPD) conditions imposed inside the GraPh platform during the phenotyping of transpiration rate (TR) response curves to increasing VPD of the NAM25 recombinant inbred lines (RILs) and their parents. For a given day, conditions are averaged across three growth chambers (3 sensors per chamber). Each VPD level is indicated with a number (1: lowest VPD; 7: highest VPD).

| Exp. | Week | Day | T (°C)<br>± S.E. | VPD1<br>(kPa)<br>± S.E. | VPD2<br>(kPa)<br>± S.E. | VPD3<br>(kPa)<br>± S.E. | VPD4<br>(kPa)<br>± S.E. | VPD5<br>(kPa)<br>± S.E. | VPD6<br>(kPa)<br>± S.E. | VPD7<br>(kPa)<br>± S.E. |
| --- | --- | --- | --- | --- | --- | --- | --- | --- | --- | --- |
| E1 | Week 1 | 1 | 30.3 ± 0.02 | 1.5 ± 0.0 | 1.9 ± 0.0 | 2.3 ± 0.0 | 2.4 ± 0.0 | 2.6 ± 0.0 | 2.8 ± 0.0 | 2.9 ± 0.0 |
|  |  | 2 | 30.3 ± 0.02 | 1.5 ± 0.0 | 1.7 ± 0.0 | 2.3 ± 0.0 | 2.5 ± 0.0 | 2.7 ± 0.0 | 2.9 ± 0.0 | 3.0 ± 0.0 |
|  |  | 3 | 30.2 ± 0.03 | 1.4 ± 0.0 | 1.7 ± 0.0 | 2.2 ± 0.0 | 2.4 ± 0.0 | 2.6 ± 0.0 | 2.8 ± 0.0 | 2.9 ± 0.0 |
|  | Week 2 | 1 | 30.4 ± 0.02 | 1.5 ± 0.0 | 1.8 ± 0.0 | 2.2 ± 0.0 | 2.4 ± 0.0 | 2.7 ± 0.0 | 2.9 ± 0.0 | 3.0 ± 0.0 |
|  |  | 2 | 30.1 ± 0.02 | 1.5 ± 0.0 | 1.8 ± 0.0 | 2.2 ± 0.0 | 2.4 ± 0.0 | 2.6 ± 0.0 | 2.8 ± 0.0 | 3.0 ± 0.0 |
|  |  | 3 | 30.3 ± 0.02 | 1.5 ± 0.0 | 1.9 ± 0.0 | 2.2 ± 0.0 | 2.5 ± 0.0 | 2.7 ± 0.0 | 3.0 ± 0.0 | 3.0 ± 0.0 |
|  | Week 3 | 1 | 30.6 ± 0.02 | 1.5 ± 0.0 | 1.9 ± 0.0 | 2.3 ± 0.0 | 2.6 ± 0.0 | 2.7 ± 0.0 | 2.9 ± 0.0 | 3.0 ± 0.0 |
|  |  | 2 | 30.3 ± 0.02 | 1.5 ± 0.0 | 1.9 ± 0.0 | 2.3 ± 0.0 | 2.5 ± 0.0 | 2.7 ± 0.0 | 2.9 ± 0.0 | 3.0 ± 0.0 |
|  |  | 3 | 30.4 ± 0.01 | 1.5 ± 0.0 | 1.9 ± 0.0 | 2.2 ± 0.0 | 2.4 ± 0.0 | 2.6 ± 0.0 | 2.9 ± 0.0 | 3.1 ± 0.0 |
| E2 | Week 1 | 1 | 29.8 ± 0.01 | 1.4 ± 0.0 | 1.8 ± 0.0 | 2.0 ± 0.0 | 2.3 ± 0.0 | 2.5 ± 0.0 | 2.8 ± 0.0 | 3.0 ± 0.0 |
|  |  | 2 | 29.9 ± 0.02 | 1.4 ± 0.0 | 1.8 ± 0.0 | 2.2 ± 0.0 | 2.4 ± 0.0 | 2.6 ± 0.0 | 2.9 ± 0.0 | 3.0 ± 0.0 |
|  |  | 3 | 30.1 ± 0.03 | 1.5 ± 0.0 | 1.8 ± 0.0 | 2.3 ± 0.0 | 2.4 ± 0.0 | 2.6 ± 0.0 | 2.9 ± 0.0 | 3.1 ± 0.0 |
|  | Week 2 | 1 | 29.9 ± 0.01 | 1.5 ± 0.0 | 1.9 ± 0.0 | 2.1 ± 0.0 | 2.4 ± 0.0 | 2.7 ± 0.0 | 2.9 ± 0.0 | 3.0 ± 0.0 |
|  |  | 2 | 30.0 ± 0.02 | 1.5 ± 0.0 | 1.9 ± 0.0 | 2.1 ± 0.0 | 2.4 ± 0.0 | 2.7 ± 0.0 | 3.0 ± 0.0 | 3.1 ± 0.0 |
|  |  | 3 | 29.8 ± 0.01 | 1.5 ± 0.0 | 1.8 ± 0.0 | 2.1 ± 0.0 | 2.3 ± 0.0 | 2.6 ± 0.0 | 2.9 ± 0.0 | 3.1 ± 0.0 |
|  | Week 3 | 1 | 30.0 ± 0.01 | 1.5 ± 0.0 | 1.9 ± 0.0 | 2.1 ± 0.0 | 2.4 ± 0.0 | 2.7 ± 0.0 | 3.0 ± 0.0 | 3.1 ± 0.0 |
|  |  | 2 | 29.8 ± 0.01 | 1.5 ± 0.0 | 1.9 ± 0.0 | 2.1 ± 0.0 | 2.4 ± 0.0 | 2.6 ± 0.0 | 2.9 ± 0.0 | 3.0 ± 0.0 |
|  |  | 3 | 29.8 ± 0.01 | 1.4 ± 0.0 | 1.8 ± 0.0 | 2.1 ± 0.0 | 2.3 ± 0.0 | 2.6 ± 0.0 | 2.9 ± 0.0 | 3.1 ± 0.0 |

In each chamber, environmental conditions (T, RH, VPD, and PAR) at canopy level were continuously recorded every 5 min in three locations by the same sensors as described earlier.

On the day preceding the phenotyping, plants were progressively watered until dripping and left to drain for approximately 6 hours, a period during which pots were covered with aluminium foil, to nullify soil water evaporation. Thereafter, three replicates of each genotype were randomly placed on three balances across three growth chambers during the afternoon (one replication per chamber). During that time, plants were exposed to environmental conditions that are similar to the last day for a few hours, before exposing plants to uniform night-time conditions (duration: from 20:00- to 06:00; PAR = 0 µmol m^-2^ s^-1^; T = 20°C and VPD = 1.0 kPa). The phenotyping sequence started at 06:00 the next morning following the procedure of Tamang *et al*. (2022). In short, the seven target VPD settings were imposed for 60 min each, sequentially from the lowest to the highest, combining the use of industrial foggers and programmable de-humidifiers. At the end of the sequence, plants were removed from the platform and transferred to the lab where leaf areas were measured destructively using a leaf area meter (model LI3100-C, LiCOR, Lincoln, NE) so that transpiration rates account for differences in canopy size across genotypes. In total, during the two phenotyping experiments, the phenotyping platform enabled screening 900 plants.

### Data analysis

#### Fitting transpiration rate response curves to increasing vapor pressure deficit

Whole plant transpiration rate (TR, mol H_2_O m^2^ s^-1^) was estimated gravimetrically and averaged every 15 minutes within each VPD step using the following equation:

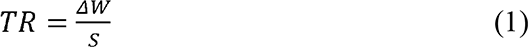

Where Δ*W* is the change in weight over time recorded by the balances and *S* is the whole-plant one-sided leaf surface area.

For each individual plant, values for TR were computed for each VPD step following Tamang and Sadok (2018). Three average TR and VPD values were calculated over a period of 45 mins (15 minutes each) for each 60 min step, as plants are likely to be acclimating to the new VPD regime during the first 15 min (Fletcher *et al*. 2007; Schoppach and Sadok 2012). In total, 42 data points were obtained for each genotype (2 experiments X 7 VPD steps X 3 replicate plants) in the 1.5–3 kPa VPD range.

In order to characterize the diversity of response curves at the individual plant level, TR vs VPD regressions were first subjected to two fits: one linear and one segmented following Tamang *et al*. (2019). Briefly, the two regression models, were compared for each individual plant and the best fitting formalism was determined based on an extra sum-of-squares F test (*P* < 0.05). This analysis revealed that a first group of plants were better described by a linear formalism while a segmented one applied best to the rest, and that for certain genotypes, individual replicates exhibited both response types. This made it unsuitable to use this bi-modal fitting approach for phenotyping purposes due to the need for extracting the same phenotypic data from all genotypes to conduct the genetic mapping. Therefore, fitting TR vs VPD response curves was carried out at the individual plant level using a quadratic model (Monteith 1995) for all genotypes, as follows:

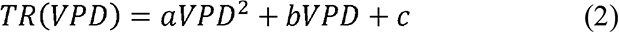

Where the parameters *a* and *b* are the quadratic and linear terms of the model, respectively, and *c* is the y intercept. Using this equation, TR at 1.5 and 3.0 kPa were estimated for each individual replicate plant, i.e., *TR_1.5_* and *TR_3.0_* (Fig. 1a). Additionally, the maximum transpiration rate, *TR_MAX_*, was calculated as the TR obtained at the maximum of eq. 2, i.e., when TR is at the point where the first derivative of eq. 2 equals 0. The theoretical VPD expected at this point, was called VPD breakpoint for transpiration rate (*VPD_BP,TR_*). The model parameters *a*, *b*, and *c* were estimated using in R-based scripts (R Core Team 2016) including the *nls* function and the port algorithm (Bates and Watts 1988). Consistent with the assumption that the quadratic model has a maximum, the quadratic term was given lower and upper bounds of negative infinity and zero, respectively.

**Fig. 1.**
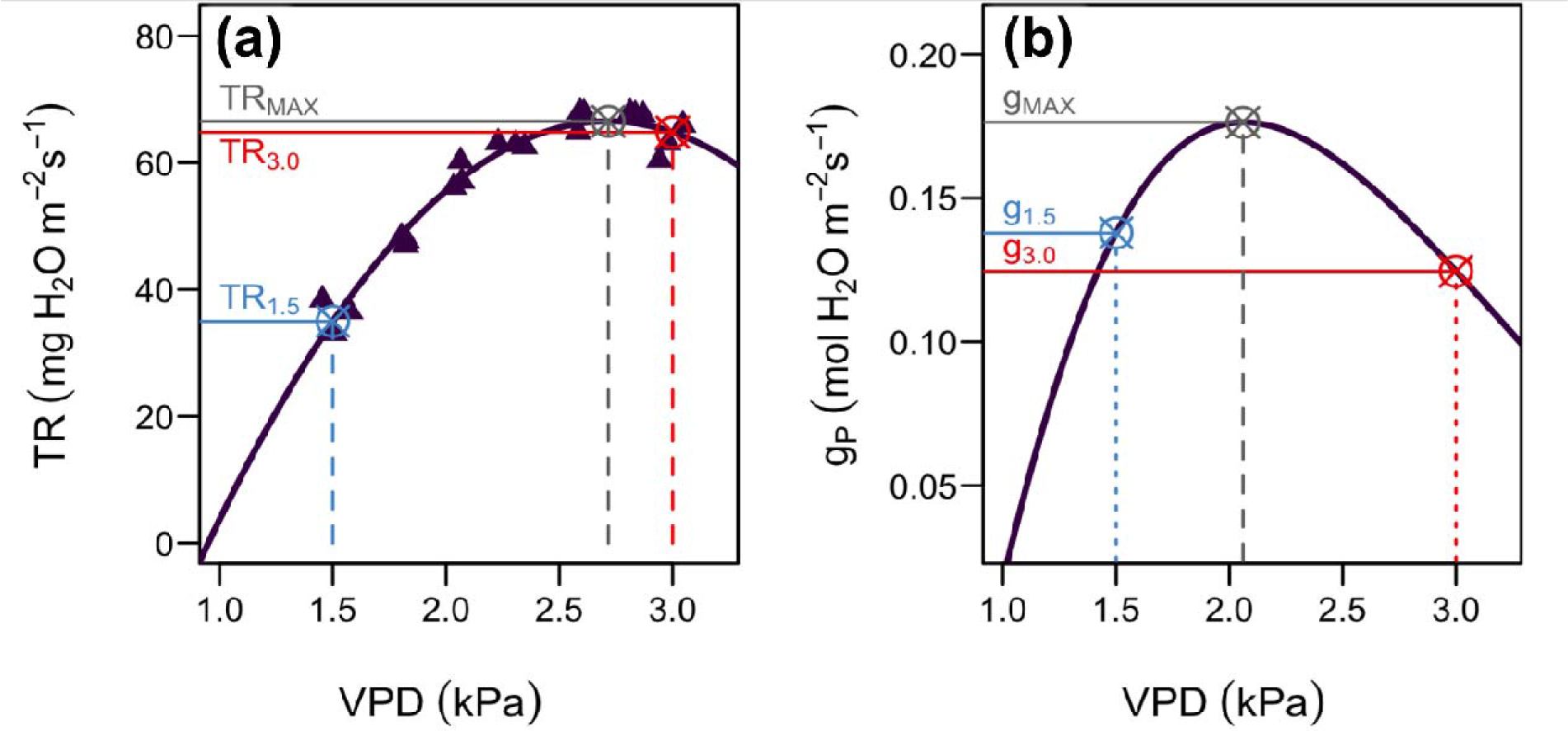
Examples illustrating how transpiration rate (TR) and whole-plant stomatal conductance (*g_P_*) were derived for each individual plant based on response curves to increasing vapor pressure deficit (VPD). The curves represent the response of a replicate plant from RIL 2 measured during experiment 2. In both panels, open crossed circles represent calculated values from the curve, vertical dashed lines indicate the corresponding VPD values and horizontal solid lines indicate the corresponding transpiration rate (TR, panel A) or whole-plant conductance (*g_P_*, panel B). In panel A, TR estimated at 1.5 kPa (*TR_1.5_*), and 3 kPa (*TR_3.0_*), and the maximum TR (*TR_MAX_*) are coloured light blue, red and grey, respectively. In panel B, the corresponding conductance parameters *g_PMAX_*, *g_P1.5_*, and *g_P3.0_* are reported following an identical colour scheme.

Subsequently, whole-plant stomatal conductance (*g_P_*) was calculated using the equation derived in Sadok *et al*. (2020):

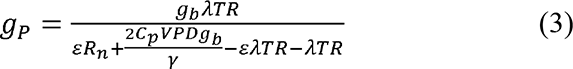

Where A is molar latent heat of vaporization of water, ε is the ratio of the slope of the relation between saturation vapor pressure and T to γ (the psychrometric constant), *R_n_* is net radiation flux density, *C_p_*is the molar heat capacity of dry air at constant pressure, *g_b_*is the boundary layer surface conductance per area unit *S*. The parameters used to solve this model were those described in Sadok *et al*. (2020) except for g_b_ which was 0.4 mol H_2_O m^2^ s^-1^ instead of 0.2 because Sadok *et al*. (2020) solved the equation for each side of the leaf, and the conversion of *R_n_* was approximated as *R_n_* ≈ *PAR/4.8*, assuming a fluorescent light similar to the one reported by (Thimijan and Heins 1983).

This equation was solved for each value of TR obtained from equation 2 between 1 and 3 kPa in 0.01 kPa increments. The maximum *g_P_* value within this range was reported as *g_PMAX_*. In the very rare cases (only two) where the curve did not reach its maximum within this range, values of *g_PMAX_* for those plants were not reported. The VPD value at which *g_PMAX_* was reached was called VPD breakpoint for whole plant stomatal conductance, or *VPD_BP,gP_*(Fig. 1b).

Additionally, whole-plant stomatal conductance (*g_P_*) at 1.5 and 3.0 kPa were estimated for each plant as *g_P1.5_* and *g_P3.0_* (Fig. 1b).

#### Phenotypic values and QTL mapping

A linear model accounting for the nested experimental design was fit to the data. The main effects of RIL, experiment, week nested within experiment, day nested within week, and growth chamber nested within day were included in the model as fixed effects. Best linear unbiased estimates of RILs for each phenotype were output from the model and used for QTL mapping. For each of the traits described above, namely *TR_1.5_*, *g_P1.5_*, *TR_3.0_*, *g_P3.0_*, *TR_MAX_*, *g_PMAX_*, *VPD_BP,TR_*, *VPD_BP,gP_*, broad-sense heritability (H^2^) coefficients were estimated on an individual- pot basis using the R package ‘heritability’ (Kruijer *et al*. 2015). Mean phenotypic values were calculated using the libraries lmerTest (Kuznetsova *et al*. 2017) and lsmeans (Lenth 2016) in R (R Core Team 2016). Variables *g_P1.5_*, *g_P3.0_*, and *g_PMAX_* were log-transformed for the analysis in order to meet the normality assumption. The antilog of the results was reported when appropriate.

The NAM25 population used in this study was genotyped (Song *et al*. 2017) using the Illumina Infinium SoyNAM6K Bead Chip developed for this population made publicly available in the SoyBase data base (Grant *et al*. 2010). Out of the 4312 single nucleotide polymorphism (SNP) markers found to be polymorphic and informative for the whole population, only 1534 markers were found to be polymorphic and informative for this population. Monomorphic markers as well as markers exhibiting complete linkage disequilibrium with each other were filtered with the aid of the library *qtl* in R (Broman *et al*. 2003).

Quantitative trait loci (QTLs) were identified using the library qtl2 in R (Broman *et al*. 2019). The additive model was used to estimate marker probability and the leave one chromosome out (LOCO) method was used for the QTL analysis. The population was simulated as an F_5_ advanced intercross. The logarithm of the odds (LOD) detection threshold was approximated through a permutation test with 1000 permutations. Confidence intervals for the QTLs were approximated using the 1.5 LOD approach. The coefficient of determination (R^2^) and the additive effect of the QTLs were estimated through a linear least-squares regression of the phenotypic value on allelic dosage (Lynch and Walsh 1998).

## Results

There was a substantial variation in TR and *g_P_* response curves to increasing VPD in the studied population. This variation is exemplified in Fig. 2 which illustrates the range of TR and *g_P_* response curves to VPD among two contrasted lines (RIL 188 and RIL 121), based on all replications and experiments. Fitting regressions on all genotypes following the example illustrated on Fig. 2 yielded high R^2^ values (0.92 to 0.99 across all replicates) with parameters *TR_1.5_, TR_3.0_, TR_MAX,_ g_P1.5_, g_P3.0,_ g_PMAX_* exhibiting a substantial genetic variability and normal distributions (Fig. 3). Particularly, parameters *TR_1.5_, TR_3.0_, TR_MAX_* and their derived whole-plant stomatal conductance counterparts varied two-fold or more across genotypes. For instance, *TR_1.5_* ranged from 19 to 38 mg H_2_O m^-2^ s^-1^ (Fig. 3a), while parameter *TR_3.0_*exhibited a similar variation, although at a higher range (33 to 64 mg H_2_O m^-2^ s^-1^, Fig. 3c). The estimated VPD breakpoints, however, were more tightly grouped, averaging approx. 3.1 and 2 kPa for *VPD_BP,TR_* and *VPD_BP,gP_*, respectively (Fig. 3g,h). Frequency distributions showcased transgressive segregation for all traits (Figure 3).

**Fig. 2.**
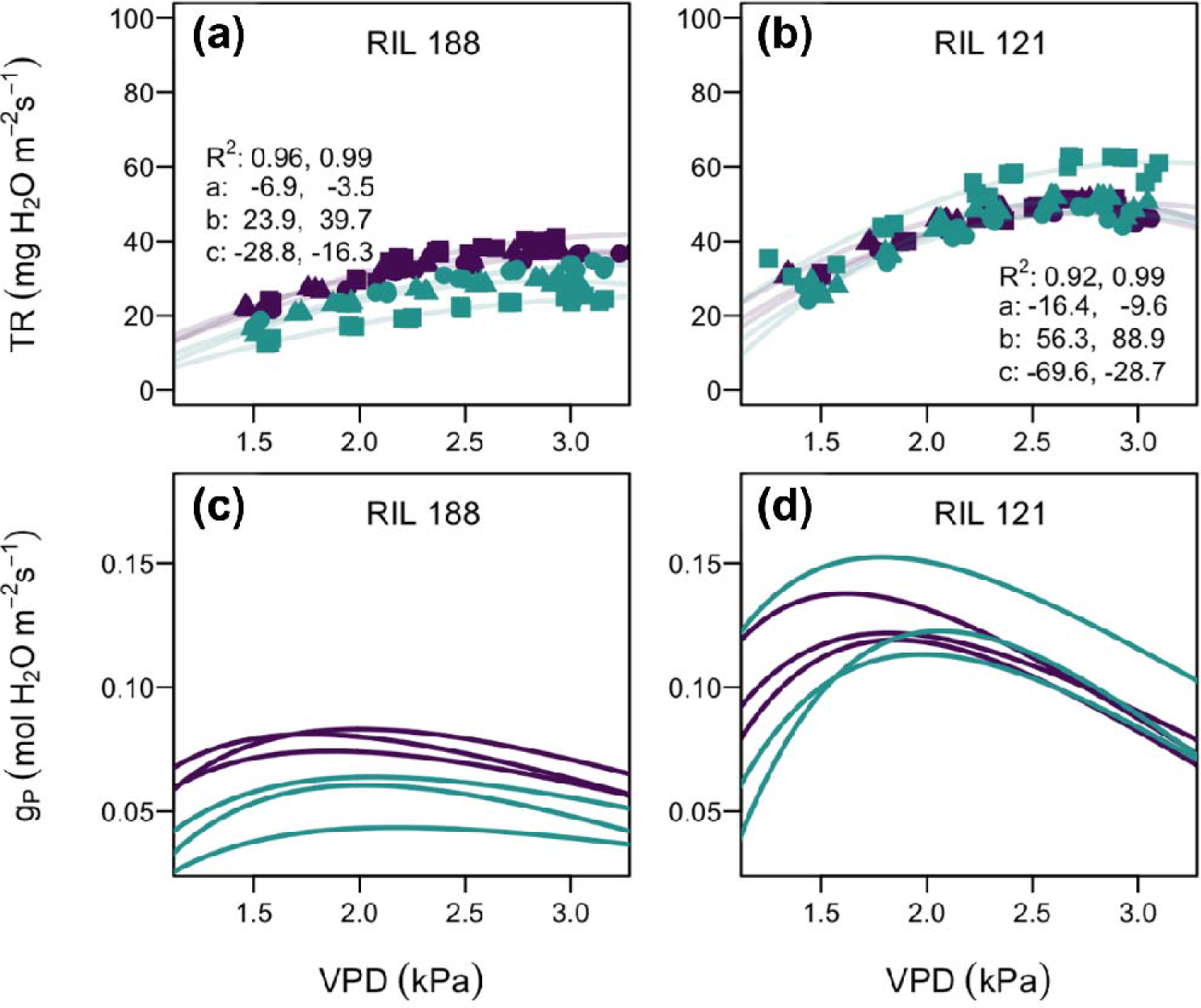
Example of response curves of transpiration rate (TR, panels a, b) and derived whole- plant stomatal conductance response curves (g_P_, panels C,D) to increased vapor pressure deficit (VPD) for two contrasted recombinant inbred lines (RILs) from the studied population. In panels A and B, the different colours and symbols represent different experiments and replicate plants, respectively. The range of parameters a, b, c, and coefficient of determination (R^2^) of TR responses to VPD across all 6 replications is reported.

**Fig. 3.**
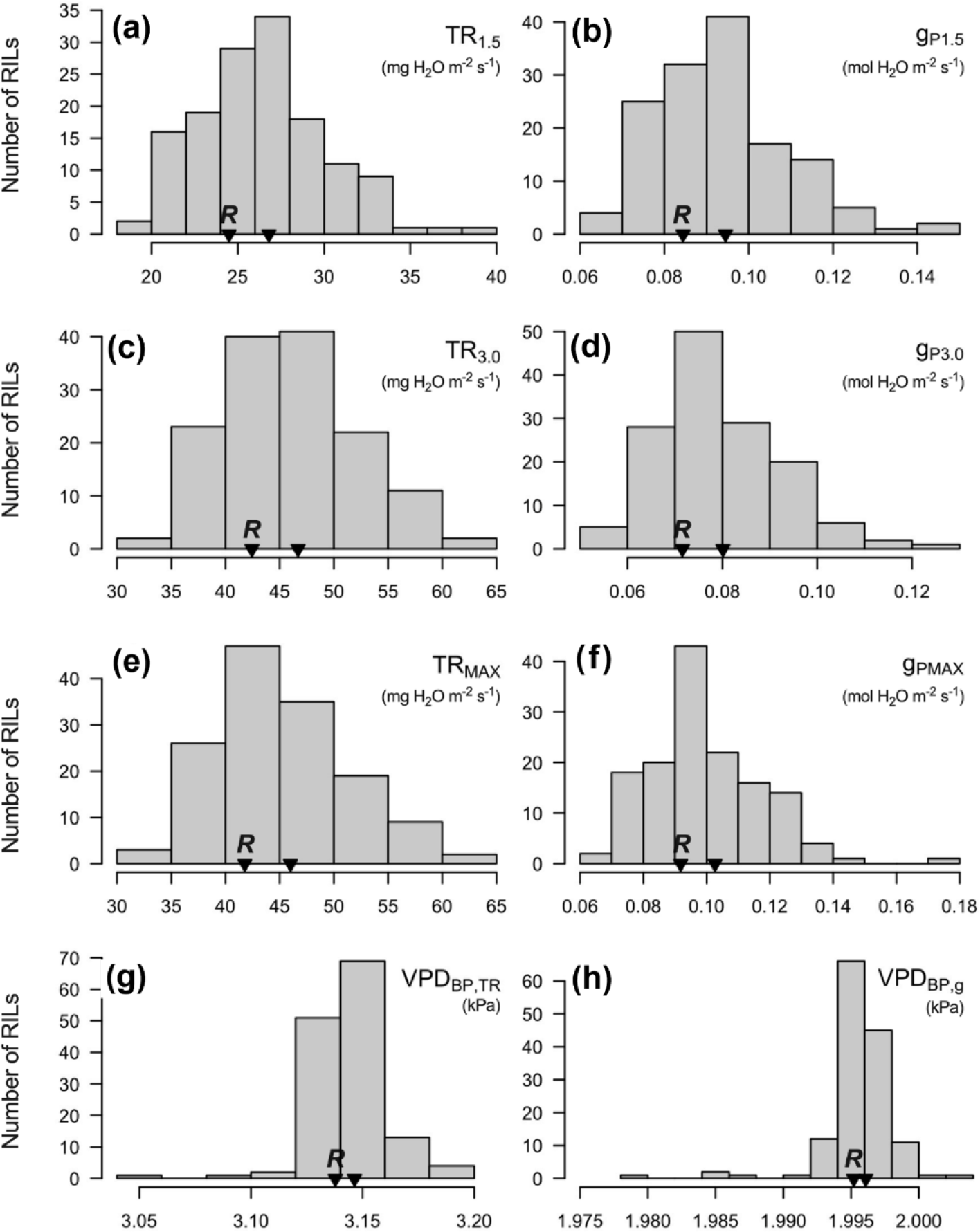
Frequency distribution of trait values for the NAM25 recombinant inbred lines (RILs) population phenotyped for transpiration rate (TR) response to increasing vapor pressure deficit (VPD). In each panel, the name of the trait, its unit, and the position of the recurrent (*R*) parent (i.e., IA3023) are indicated.

As compiled in Table 3 and illustrated in Fig. 4, this variation translated into significant QTL for all traits, except for VPD breakpoint parameters. More specifically, two QTL were detected for *TR_1.5_*, *TR_3.0_*, and *TR_MAX_* on chromosomes 6 and 10 (Table 1; Fig. 4a,c,e), while one additional transpiration QTL was found to be specific to low VPD conditions, i.e., detected only for *TR_1.5_* on chromosome 4 (Fig. 4a). Among these TR QTL, the one detected on chromosome 6 had the highest explained phenotypic variation (R^2^) which ranged from 0.10 to 0.15 depending on the trait (i.e*., TR_1.5_*, *TR_3.0_*, or *TR_MAX_*_;_ Table 1). The QTL detected for the conductance parameters (*g_P_*) generally mirrored those identified for TR, confirming the QTL detected on chromosomes 4 and 6, but were not found to be associated with the TR QTL detected on chromosome 10 (Table 1; Fig. 4b,d,f). However, similar to TR QTL, the *g_P_* QTL detected on chromosome 6 exhibited the highest phenotypic variance explained (R^2^ = 0.10 to 0.16).

**Fig. 4.**
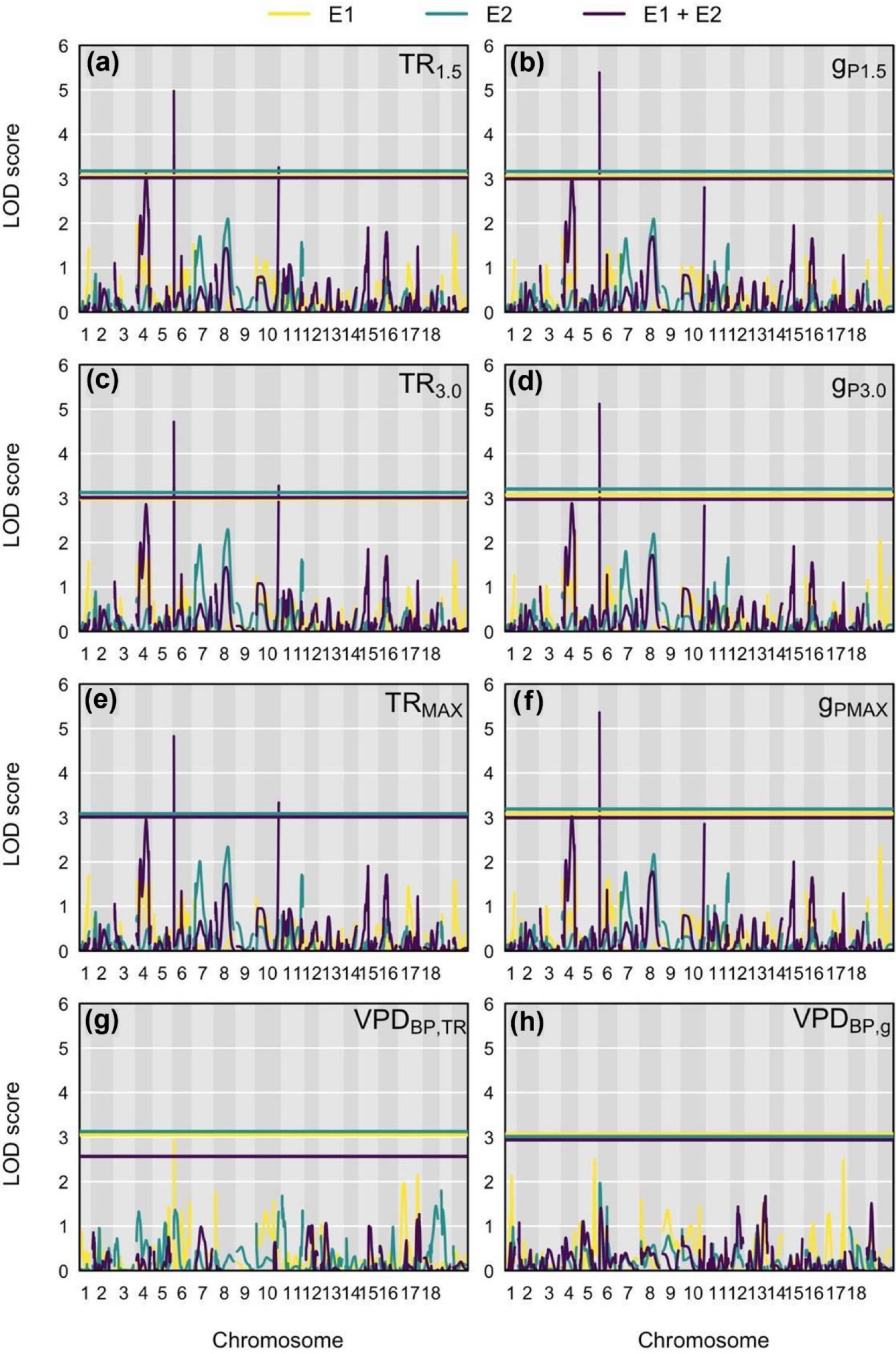
LOD score curves of the QTL mapping conducted for all examined traits as a function of the experiment. In each panel, the name of the trait is indicated, with different colours reflecting phenotypic data from experiments E1 and E2 used separately or combined in the analysis. Chromosomal positions are projected on the x-axis and horizontal lines illustrate LOD significance thresholds.

**Table 3.** Summary of significant QTLs (α = 0.05) detected for the traits describing whole-plant transpiration rate (TR) response to increasing vapor pressure deficit (VPD) in the mapping population. Abbreviations: Chr: Chromosome; Pos: Position (cM); LL: QTL lower limit (cM); UL: QTL upper limit (cM); LOD: logarithm of the odds; R^2^: coefficient of determination; Effect: Additive effect of individual alleles from parent LG03-3191.

| **Trait** | **Chr.** | **Pos** | **LL** | **UL** | **LOD** | **$R^2$** | **Effect** |
| --- | --- | --- | --- | --- | --- | --- | --- |
| TR<sub>1.5</sub> | 4 | 65.54 | 28.99 | 81.25 | 3.20 | 0.06 | -0.00006 |
|  | 6 | 0 | 0 | 0.85 | 4.97 | 0.13 | -0.00008 |
|  | 10 | 104.36 | 96.79 | 104.36 | 3.28 | 0.08 | 0.00006 |
| TR<sub>3.0</sub> | 6 | 0 | 0 | 0.85 | 4.73 | 0.10 | -0.00012 |
|  | 10 | 104.36 | 96.79 | 104.36 | 3.30 | 0.07 | 0.00010 |
| TR<sub>MAX</sub> | 6 | 0 | 0 | 0.85 | 4.89 | 0.15 | -0.00015 |
|  | 10 | 104.36 | 96.79 | 104.36 | 3.39 | 0.07 | 0.00011 |
| g<sub>P1.5</sub> | 4 | 66.34 | 28.99 | 81.25 | 3.11 | 0.06 | -0.05 |
|  | 6 | 0 | 0 | 0.85 | 5.40 | 0.16 | -0.07 |
| g<sub>P3.0</sub> | 6 | 0 | 0 | 0.85 | 5.16 | 0.10 | -0.06 |
| g<sub>PMAX</sub> | 4 | 66.34 | 28.99 | 81.25 | 3.11 | 0.08 | -0.05 |
|  | 6 | 0 | 0 | 0.85 | 5.38 | 0.14 | -0.07 |

Overall, irrespective of the trait, the QTL on chromosome 6 was detected for all 6 traits, spanned a much smaller genetic distance (0.85 cM) compared to all other QTL which covered much larger genetic distances (3 to 4 orders of magnitude bigger). The chromosome 6 QTL also consistently exhibited the highest LOD scores and the strongest additive effects. In all instances where this QTL was detected, the allele from LG03-3191 consistently conferred a negative additive effect (Table 1).

## Discussion

Currently, very little information is available on the genetic basis of whole-plant TR response to increasing VPD in plants, with only one study so far documenting a genetic basis for such response curves conducted on wheat (Tamang *et al*. 2022). Despite a relatively large body of research dedicated to screening TR response curves to VPD in soybean (e.g., Fletcher *et al*. 2007; Sinclair *et al*. 2008; Sadok and Sinclair 2009ab; Seversike *et al*. 2014; Devi *et al*. 2014), no QTL have been detected for traits directly reflecting such response curves in soybean.

The QTL mapping performed here on the NAM25 population resulted in the detection, for the first time, of a VPD-dependent QTL controlling whole-plant TR, found on chromosome 4 for *TR_1.5_* but not for *TR_3.0_*or *TR_MAX_* (Table 3). This result is consistent with a similar finding on wheat where a single nucleotide polymorphism (SNP) controlling *TR_1.5_* was identified in a region that was unique to this trait (Tamang *et al*. 2022). Furthermore, this study revealed a potentially robust QTL on chromosome 6, which was detected for all 6 transpiration and conductance traits, spanning the smallest genetic distance (0.85 cM) and explaining the highest level of phenotypic variance among all identified loci (Fig. 4; Table 3). While such QTL will need to be confirmed in future studies, our results indicate that TR response to VPD is under the control of genetic loci located on chromosomes 4, 6, and 10 in soybean.

The detection power of our analysis was relatively low, shown by heritability values around 0.1, so the fact we still were able to detect QTL indicates that large-effect QTL do likely exist. In any case, confirmed QTL could serve as a basis for marker-assisted breeding to design cultivars adapted to contrasted water availability regimes. For instance, cultivars harbouring QTL alleles conferring reducing TR at high VPD would be prime targets for deployment in environments where terminal drought occurs, since TR sensitivity to VPD has been shown to lead to higher yields through water conservation (i.e., Fletcher *et al*. 2007; Sinclair *et al*. 2010; Ye *et al*. 2020). In contrast, ‘water-consumptive’ genotypes with alleles maximizing TR or *g_P_* regardless of the VPD range would be much better suited to environments where soil moisture is not -or rarely- limiting. In this case, such alleles would be favourable because they maximize nutrient and water uptake, both of which are critical for canopy growth and yield (Sinclair *et al*. 2017; Sadok *et al*. 2019). Increased expenditure under high VPD could be also highly beneficial in environments where water is not limiting where transpirational cooling protects from leaves and developing pods from heat stress (Sadok *et al*. 2021).

The relatively limited number of QTL detected in this study could be explained by three main factors. One consists of the need for a sequential screening approach. Indeed, concomitantly phenotyping the entire mapping population implies tracking whole-plant water use on more than 450 plants using balances, an approach that is currently extremely cost-prohibitive. As a result, our experiment necessitated three weekly sequential plantings, which introduced the potential for developmental differences between the three planting groups as greenhouse conditions will not be exactly uniform across groups. Another important factor to consider is the genetic structure of the mapping population. This population was not designed based on parents that segregate strongly for the trait of interest, but rather, was chosen from a pool of pre-existing NAM populations. Perhaps QTL could be found in greater resolution or quantity if a population was constructed from two highly contrasting parental varieties which could create more diverse progeny for QTL mapping, although that requires a much greater effort including breeding, genotyping, and linkage map construction. Finally, our phenotyping approach focused solely on varying VPD, in order to reduce interactions with undesired, typically co-varying sources of environmental variation such as light, temperature, and wind, which non-linearly impact TR (Sadok *et al*. 2021). This phenotyping strategy might have contributed to reducing the number of QTL found. Thus, a follow-up to this investigation would be to confirm presence of the identified QTL in a different genetic background that maximizes potential allelic diversity for this trait, using a phenotyping approach that minimizes the need for sequential planting.

Overall, this study demonstrated for the first time that plant transpirational water loss response to atmospheric drying, a trait that has been associated with drought tolerance in soybean, has a tractable genetic basis. The results open a new, untapped avenue for identifying alleles enabling water conservation under water-limited environments or for higher water use in environments where such a strategy is beneficial.

## Conflicts of interest

The authors declare no conflicts of interest.

## Declaration of funding

This work was supported by USDA NIFA through the Minnesota Agricultural Experiment Station (project# MIN-13-095), the Minnesota Soybean Research & Promotion Council (Projects# 00063035, 00070622 and 00078080). Support from the Jean W. and Mary S. Lambert Agronomy and Plant Genetics Fellowship is gratefully acknowledged. The authors declare no conflicts of interest.

## Author contributions

WS and AJL conceived the study; DM, BT, EM conducted the experiments; JL and DM performed the data analysis. DM, JL and WS wrote the first draft of the manuscript.

## Data availability

The data that support this study will be shared upon reasonable request to the corresponding author.

## ORCID

Walid Sadok https://orcid.org/0000-0001-9637-2412 Aaron J Lorenz https://orcid.org/0000-0002-4361-1683

## Notes

### Competing Interest Statement

The authors have declared no competing interest.

